# Dissociated responses of vesiculogenesis and amoxicillin impact on extracellular vesicle production of first gut bacterial colonizers *Bifidobacterium longum* and *Lactiplantibacillus plantarum*

**DOI:** 10.64898/2026.07.27.740896

**Authors:** Anaïs Halbert, Amandine Dupuy, Lisa Wallart, Sophie Brouard, Julie Hardouin, Hervé M. Blottière, Odile Tresse

## Abstract

Bacterial extracellular vesicles (bEVs) have emerged as important mediators of microbiota–host interplay through the transport of active biomolecules, namely cargos, far from their release location. The neonatal period represents a critical window for the establishment of the gut microbiota and subsequent sustainable symbiotic communication. The gut primo-colonizing bacteria, including bifidobacteria and lactobacilli, likely contribute to the impact on the digestive, immune and neuron system maturation. However, exposures and experiences during this early stage may influence the development of health and diseases later on in life by altering these primo-interactions. As antibiotherapies are frequent in the postnatal period and associated to microbiota disorders, we evaluated the impact of amoxicillin on first colonizing Gram positive-derived EVs, using a robust and reproductible in-house workflow for the extraction and purification bEVs from *Bifidobacterium longum* and *Lactiplantibacillus plantarum*. The EVs production and the proteovesiculome profiles under amoxicillin treatment were compared. The results pointed out a dissociated response in the EVs release process and their regulation by amoxicillin according to strain with an enhance production of EVs for *B. longum* under amoxicillin. In addition, the proteovesicular analyses indicate that the vesicular protein profile was enriched and more diverse in *B. longum*-derived EVs from amoxicillin-treated cells than those from non-treated cells while the content shift in *L. plantarum*-derived EVs in amoxicillin-treated cells was in favor of protein richness loss. Overall, this study opens new avenues considering the impact of antibiotic therapies in the neonatal period on EVs derived from benefit Gram-positive gut bacteria.

## INTRODUCTION

The Developmental Origins of Health and Disease (DOHaD) is a concept that emphasizes the impact of early life, particularly the first thousand days, from conception to a child’s second birthday, on the development of health across the lifespan (Hanson and Gluckman, 2014). Over the two last decades, the gut microbiome was identified as a key factor in metabolic programming that could impact the individual’s susceptibility to onset metabolic disorders (Catassi et al., 2024). Gut colonization occurs after birth with a diversity among micro-organisms depending on various factors including -but not limited to-genetics, gestational age, mode of delivery, mode of feeding and environment (Laforest-Lapointe and Arrieta, 2017; Roswall et al., 2021). At birth, the newborn acquires approximately 50% of its microbial species directly from the mother, mainly through the intestinal tract, vagina, skin, and oral cavity. Among the bacteria transmitted from mother to child, the most common belong to the phyla Actinomycetota and Bacillota, including Bifidobacteriaceae and Lactobacilliaceae (Solís et al., 2010; Bäckhed et al., 2015; Ferretti et al., 2018, Mancabelli et al., 2020). The interplay between these first colonizing bacteria and the host cells is highly dynamic and it underscores the importance of a balanced microbial ecosystem, where both the host and its microbial residents benefit from this balance to prevent disease developments. However, the initiation of the interplay by the first commensals remains poorly described, moreover when it is constantly challenged by traumatic episodes (van de Guchte et al., 2018).

Antibiotic exposure is one of the widespread extrinsic factors influencing the shaping and fitness of gut microbiota after birth as bacterial infections are more frequent during this period. Indeed, many newborns are treated with antibiotic, most often in response to suspected early-onset neonatal infection or maternal risk factors during delivery. These antibiotic therapies are based on broad-spectrum penicillins, with amoxicillin (AMX) being the most frequently prescribed (CDC, 2024). Even short-term antibiotic treatment can exert long-lasting effects on the gut microbiota (Jernberg et al., 2010). Several studies have shown that early antibiotic exposure is often associated with an increased abundance of the phyla Bacteroidota and Pseudomonodota, alongside a decrease in Actinomycetota and Bacillota (Fouhy et al., 2012; Korpela et al., 2016; Tapiainen et al., 2019). In addition, antibiotic administration during the first days of life leads to a reduction in bacterial diversity (Tanaka et al., 2009). Although some of these alterations may be transient and microbial composition may recover after discontinuation of treatment, the metabolic functions associated with the microbiota can remain disrupted over the long term (Cox et al., 2014; Korpela et al., 2016). Several cohort studies have reported associations between repeated or early-life antibiotic exposure and an increased risk of childhood obesity, asthma, eczema, and allergic diseases (Aversa et al., 2021). These findings highlight the long-term impact of early microbiota disturbances and emphasize the importance of research on antibiotic use during pregnancy and early childhood.

The predominance of bifidobacteria, particularly *Bifidobacterium longum,* in the infant gut is due to their ability to metabolize human milk oligosaccharides (HMOs) for their growth, producing meanwhile metabolites as short chain fatty acids (SCFA) like butyrate and indole derivatives like indole-3 lactate (ILA) which both contribute to the body interplay (Laursen et al., 2021; Gutierrez et al., 2023). The crosstalk of lactobacilli, specially *Lactiplantibacillus plantarum*, with the human host has been a trend scientific research concern. Properties of *L. plantarum* encompass antioxidant, antimicrobial, acidic tolerance and intestinal mucus adhesion (Echegaray et al., 2023).

Among the various mediators of communication, bacterial extracellular vesicles (bEVs) secreted by *B. longum* and *L. plantarum* have attracted growing interest due to their multifaceted biological effects and potential therapeutic applications (Abraham et al., 2025). As part of the dialogue of the primo-colonizing bacteria contributing to system maturation in newborns, bEVs transporting bioactive molecules, namely cargos, might contribute to this multifaceted and dynamic communication that underscores the far-reaching effects of gut microbes on the entire body. Although Gram-negative-derived EVs have been describing for several decades, the discovery and the studies on mEVs are more recent, overturning the paradigm that only Gram-negative bacteria could produce bEVs. The origin and the vesicular composition infer specificities which delineate the nature and the function of these bEVs. *B. longum*-derived EVs could modulate immune responses, notably by inducing mast cell apoptosis, thereby attenuating allergic reactions (Kim et al., 2016). They also promote bifidobacterial colonization and enhance adhesion to the intestinal mucosa (Nishiyama et al., 2020). Moreover, the induction of IL-10 production by extracellular vesicles from *B. longum* contributes to an anti-inflammatory immune response (Mandelbaum et al., 2023). Therapeutically, they have been shown to restore drug sensitivity in resistant ovarian cancer cells and to protect hepatocytes from fibrosis, apoptosis, and oxidative stress (Fan et al., 2024; Li et al., 2024). Altogether, these findings highlight the multifunctional role of *B. longum-*derived EVs in regulating host-microbiota interactions. *L. plantarum*-derived EVs could regulate gut microbiota composition, and attenuate inflammatory responses thereby enhancing intestinal barrier function in models of ulcerative colitis (Hao et al., 2021; Kurata et al., 2022; Chen et al., 2024; Gong et al., 2024; Tao et al., 2025). All together, these findings underscore the role of bEVs as bioactive mediators capable of influencing multiple organ systems.

In this study, by comparing *L. plantarum* and *B. longum* type-strains derived EVs, dissociated responses between the two strains were observed in the vesiculogenesis and the EVs production modulation under AMX-treated conditions. Moreover, proteovesiculomic profilings revealed that the protein content of EVs varied differently between the two strains after AMX treatment at sub-inhibitory concentrations.

## MATERIALS and METHODS

### 2.1. Bacterial strains and culture conditions

*Bifidobacterium longum* subsp *longum* JCM 1217^T^ and *Lactiplantibacillus plantarum* ATCC 14917^T^ were purchased at the Pasteur Institute Collection CIP (France) and conserved at −80°C in 20% glycerol. Strains were cultivated on Man, Rogosa, and Sharpe (MRS) agar and in MRS broth (Millipore, Sigma-Aldrich), both supplemented with 0.1% (v/v) Tween 80 (Sigma-Aldrich) and 0.5 mg/mL L-cysteine (Sigma-Aldrich) (MRS-TlC), and incubated at 37°C under anaerobic conditions using. Solid precultures were incubated at 37 °C in anaerobic conditions using the GazPak™ EZ system. Precultures were prepared in MRS-TlC and incubated at 37 °C under anaerobic conditions.

### 2.2. bEVs extraction and purification

EVs from *L. plantarum* (Lp-EVs) and *B. longum* (Bl-EVs) were isolated from culture medium incubated at 37°C for 24 h under anaerobic conditions. The bacterial growth was determined by optical density at 600 nm (OD_600_) and colony enumeration using the Miles and Misra method (Miles et al., 1938). Each culture was centrifuged at 6,000 *g* for 15 min at 4°C, followed by a second centrifugation at 8,000 *g* for 15 min at 4°C to remove bacterial cells. The culture supernatant was filtered using a 0.22 µm Stericup™ vacuum filtration system (Millipore, Sigma-Aldrich). The absence of bacterial contamination was checked by plating the filtered supernatant on MRS agar. Culture supernatant was stored at 4°C until further processing. For bEVs production under antibiotic-induced stress conditions, AMX was added to the culture medium at a sub-inhibitory concentration of 0.5 µg/mL for *B. longum* and 0.0625 µg/mL for *L. plantarum*. Sub-inhibitory concentrations of AMX were determined from the minimum inhibitory concentration (MIC) for each strain. Each condition was performed in three independent replicates.

The bEVs were then concentrated by ultrafiltration (UF) using protein concentrators with a 100 kDa molecular weight cutoff (Pierce™ PES protein concentrators, Thermo Scientific™, France). Culture supernatant was gradually loaded into the concentrators and centrifuged at 5,000 *g* at 4°C. The filter membranes were rinsed with phosphate-buffered saline (PBS) and the washed solutions were pooled. The retentate was then ultracentrifuged at 100,000 *g* for 120 min at 4°C. The pellet resuspended in PBS was submitted to size exclusion chromatography (SEC) using the qEV original 70 nm SEC column (IZON, France) following the manufacturer’s instructions. The successive eluted fractions were collected and stored at −80°C until further use. For proteomic analyses, specific aliquots containing mEVs were stored separately with Complete™ protease inhibitor cocktail Roche).

### 2.3. Protein quantification

Protein equivalent concentration of SEC fractions was determined using the Qubit™ Protein Assay Kit (Thermo Fisher Scientific). The Qubit™ working solution was prepared by diluting the Qubit™ Protein Reagent 1:200 in Qubit™ Protein Buffer. A volume of 198 µL of the working solution was added to 2 µL of each bEVs fractions (200 µL final volume) and incubated at room temperature for 15 min. Fluorescence was then measured using the Qubit™ 4 Fluorometer based a standard curve generated with known protein standards as recommended by the manufacturer.

### 2.4. Fluorescence Microscopy Analysis

Bacterial suspensions were centrifuged twice at 10,000 *g*. Then, the pellet was resuspended in PBS and fixed onto glass slides. Dual fluorescent staining was performed directly on the fixed bacteria using either CellMask™ Orange (CMO)/ 4′,6-diamidino-2-phenylindole (DAPI) or 1,1-Dioctadecyl-3,3,3,3-tetramethylindocarbocyanine perchlorate (DiI)/DAPI staining. A CMO or DiI solution was first applied onto the glass slides followed the DAPI solution according to manufacturer’s instructions. A PBS control slide was used to assess background autofluorescence. All slides were observed using a Zeiss Axio Imager epifluorescence microscope. Fluorescence signals were acquired using appropriate filter sets corresponding to the excitation and emission spectra of each dye: DAPI (excitation at 358 nm, emission at 461 nm), CMO (excitation at 554 nm, emission at 567 nm), and DiI (excitation at 549 nm, emission at 565 nm), both visualized using Rhodamine or TRITC filter sets.

### 2.5. Physical characterization of bEVs

The concentration and size distribution of particles in collected fractions after SEC were measured using nanoparticles tracking analysis (NTA) using the NanoSight NS300 device (Malvern Panalytical) equipped with a sCMOS camera. The constant flow was maintained using a syringe pump set at a speed of 50, with the chamber temperature stabilized at 25°C. For each measurement, 5 x 60 s videos were recorded with camera level 15. Data were analyzed using the NTA 3.3 Dev Build software with appropriate parameters adjusted accordingly to optimize image quality and particle detection. NTA were also performed to assess CMO-membrane labeled bEVs using the green laser (532 nm) and the 565 nm long-pass filter. The measurements obtained in the scatter mode (no filter) and the one obtained in the fluorescence mode using 565 nm long-pass filter were then compared.

### 2.6. High-Resolution Transmission Electron Microscopy

High-resolution transmission electron microscopy (HR-TEM) was used to visualize the formation of EVs on the surface of bacteria. Five µL of bacterial suspension was applied onto a Formvar/carbon-coated copper glow-discharged grids. The bacteria were then fixed using glutaraldehyde and negatively stained using UranyLess (EM-grade, Delta Microscopies, France). After air-drying, bacteria and EVs were imaged using a Themis Z G3 transmission electron microscope (Thermo Fisher Scientific) operated at 80 kV with a OneView CMOS camera (4096*4096 pixels²) in Digital Micrograph (v3.4) software (Gatan)2.8. After validated the cell distribution on the grid, half of the grid was explored and at least 15 images were taken from three different grid mesh sectors for each biological replicate.

### 2.7. Proteovesiculomic analyses

A quantity of bEVs corresponding to 25 µg of protein equivalent was mixed with 5 µL of lysis buffer (Tris-HCl pH 7.2, glycine 400 mM, 1% lauryl sarcosinate, 10 mM EDTA) and incubated for 1 h at 37 °C. After lysis, protein concentration was determined using the Qubit™ Protein Assay Kit (Thermo Fisher Scientific). For each biological replicate, 26 µg of proteins were mixed with sodium dodecyl sulfate (SDS) loading buffer (62 mM Tris-HCl pH 6.8, 20% glycerol (v/v), 0.04% bromophenol blue (w/v), 0.1 M DTT, SDS 4% (w/v)) and loaded onto a SDS-PAGE stacking gel 7%. A short electrophoresis was performed (10 mA, 15 min) to concentrate proteins. After migration, gels were stained with Coomassie Blue G250 and destained with 50% ethanol (v/v) and 10% acetic acid (v/v). The revealed protein band was excised and washed three times with water. Disulfide bonds were reduced using DTT (5 mM, 1 h) and cysteines were alkylated (20 mM iodoacetamide, 45 min in the dark). Proteins were digested with trypsin (1/26 – trypsin/protein ratio), overnight at 37 °C, in ammonium bicarbonate 10 mM, pH 8. Peptides were extracted with 100% ACN (3 times, 10 min) and then dried. Peptides were analyzed using an Orbitrap Eclipse Tribrid mass spectrometer coupled to an EASY-nLC 1 200 (both from Thermo Scientific). Peptides were solubilized in FA 1% and were injected onto an enrichment column (Acclaim PepMap100, Thermo Scientific). The separation was performed with an analytical column (Acclaim PepMap, 75µm x 25 cm C18, 2 µm, 100Å, Thermo Scientific). The mobile phase consisted of H2O/0.1% formic acid (FA) (buffer A) and CH3CN/FA 0.1% (80/20 - buffer B). Peptides were eluted at a flow rate of 300 nL/min from 2 to 40% B over 108 min. The mass spectrometer was operated in positive ionization mode with capillary voltage and source temperature set at 1.9 kV and 275 °C, respectively. The fragmentation mode was HCD (higher-energy collisional dissociation) with a collision energy of 30 eV. The first scan (MS spectra) was recorded in the Orbitrap analyzer (R = 120,000) with the mass range m/z 400–1800. Then, a cycle time of 3s was used for MS2 experiments in the orbitrap (R = 15,000). Singly charged species were excluded for MS2 experiments. Dynamic exclusion of already fragmented precursor ions was applied for 60 s and an exclusion mass width of ± 10 ppm. All measurements in the Orbitrap analyzer were performed with on-the-fly internal recalibration (lock mass) at m/z 445.12003 (polydimethylcyclosiloxane). Raw data files were processed using Proteome Discoverer 2.4 software (Thermo Scientific). Peak lists were searched using the MASCOT search software (Matrix Science) against *Bifidobacterium longum* subsp. *longum* JCM 1217^T^ databasis (ENA, AN AP010888) and the *Lactiplantibacillus plantarum* subsp. *plantarum* ATCC 14917 ^T^ databasis (ENA, AN GCA_000143745.1). Database searches were performed with the following parameters: two missed trypsin cleavage sites allowed; variable modifications: carbamidomethylation on cysteine, and oxidation on methionine. The parent-ion and daughter-ion tolerances were 5 ppm and 0.02 Da, respectively. False discovery rate (FDR) threshold for identifications was specified at 1% (for proteins and peptides). All validated protein identifications are presented in Table A1

### 2.8. Protein localization and function

For each protein sequence, the reciprocal best hit method was used to obtain the corresponding unique accession identifier in UniProt (i.e., 6 or 10 alphanumeric characters). This allowed the retrieval of Gene Ontology (GO) functional categories and canonical pathways (KEGG) using R/RStudio modules for proteomics data analyses (Tab. A2). Protein cellular localization was obtained using PSORTb-III (v3.0.3) (https://psort.org/psortb/) (Tab. A2).

### 2.9. Statistical analysis

Statistical analyses were performed using GraphPad Prism (v.10.6.1). Significant differences between groups were assessed using either Student’s t-test or one-way analysis of variance (ANOVA), as appropriate. A *p*-value < 0.05 was considered statistically significant.

## RESULTS

### 3.1. Isolation and characterization of bEVs

Without available standardized methods, isolation of bEVs is challenging in order to obtain reliable and reproducible results. As a matter of fact, protocols have to be adapted to the type of sample, the volume and the bacterial species. To address this issue, a three-step in-house protocol was developed for the extraction and purification of bEVs, including concentration by ultrafiltration followed by purification using ultracentrifugation and SEC. Protein quantification of SEC fractions (Fn) for nanoparticles derived from *B. longum* or *L. plantarum,* respectively, revealed that fractions F2 (142.8 µg/mL and 136.3 µg/mL) and F3 (117.9 µg/mL and 166.9 µg/mL) had a higher signal than fraction F1 and F4, suggesting vesicle enrichment in fractions F2 and F3 (Figs. 1A & 1C). Nanoparticle concentrations in F2 (2.11 × 10¹¹ particles/mL) and F3 (2.80 × 10¹¹ particles/mL) from *B. longum*, with an average diameter of 127.3 nm in F2 and 119.7 nm in F3 (Fig. 1B) were not significantly different, indicating the measurement reproducibility and the sample homogeneity among the three independent biological replicates. No significant differences were neither observed in concentrations (2.03 × 10¹¹ nanoparticles/mL for F2 and 2.70 × 10¹¹ nanoparticles/mL for F3) and mean sizes (133.67 nm for F2 and 139.43 nm for F3) for nanoparticles derived from *L. plantarum* (Fig. 1D). Consequently, F2 and F3 were pooled for each biological independent extraction for both strains for further analyses.

**Figure 1.**
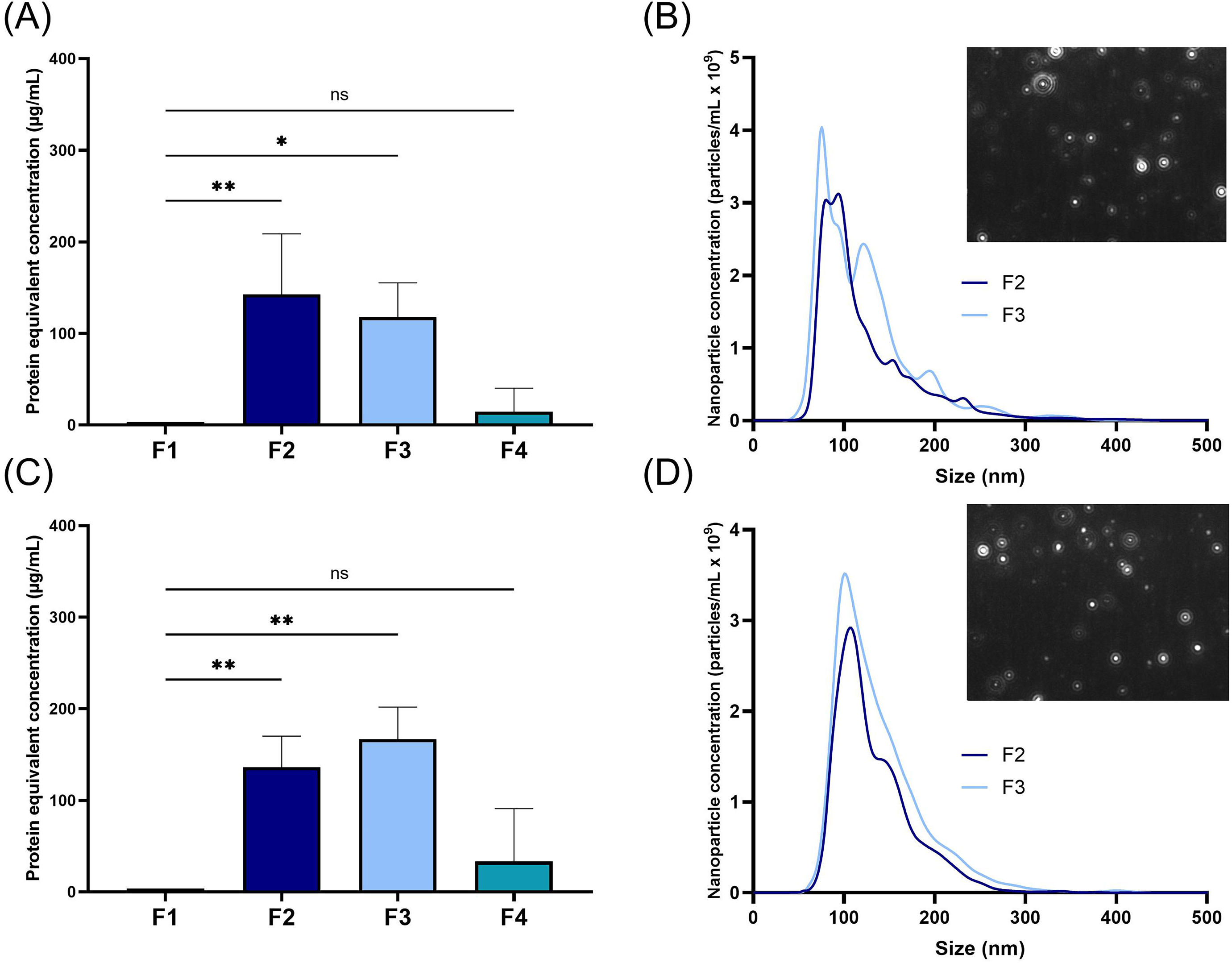
Extracted and purified nanoparticules from *B. longum* and *L. plantarum*. Analyses of protein-equivalent concentrations using Qubit quantification in purified fractions (Fn) of SEC for particles derived from *B. longum* (A) and *L. plantarum* (B). Mean of 3 biological replicates of nanoparticle concentrations according to their size with the corresponding NanoSight NS300 image for particles derived from *B. longum* (C) and *L. plantarum* (D). Measurements were performed using the optimal particle/frame rate, i.e. 20–100 particles/frame and five 1-min videos. Statistical significance is indicated as \**p* < 0.05, \*\**p* < 0.01 according to the One-way ANOVA test.

### 3.2. CellMask orange detects specifically mEVs membrane

While NTA allows quantification and size profiling of nanoparticles, it hardly differentiates mEVs from other particles. To check the efficacy of cellular membrane labeling, the strains were incubated with the DiI and the CMO in combination with DAPI to visualize cells through DNA-binding fluorescence. Both DiL and CMO dyes were able to stain *B. longum* cells, however not all the cells of *L. plantarum* were labeled using DiI while there were using CMO (Fig. 2). DiI seems to be population selective with *L. plantarum* membrane staining but not with *B. longum.* As EVs come from the cytoplasmic membrane in Gram-positive bacteria, CMO was selected for Lp-EVs and Bl-EVs labeling. NTA revealed the presence of fluorescence nanoparticles, indicating the successful integration of the dye into the vesicle membranes. As the curves from light scatter and 532-laser measurements have a similar trend for nanoparticle concentration and size, our nanoparticles measured using NTA were then assumed to be mEVs (Fig. 3).

**Figure 2.**
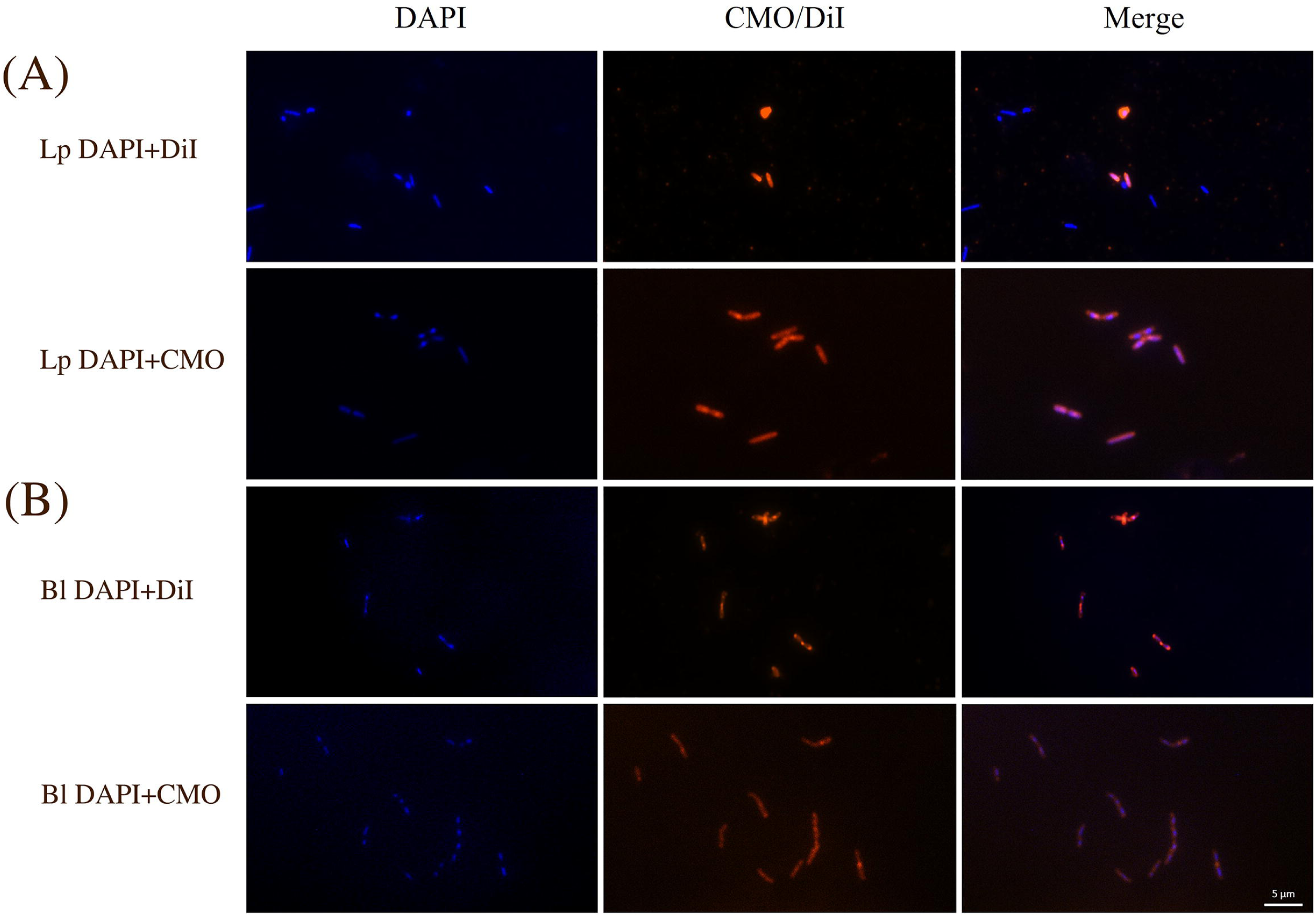
Epifluorescence imaging of *B. longum* and *L. plantarum* stained using Dil and CMO fluorophores. *B. longum* (A) and *L. plantarum* (B) stained with DAPI (left), with the DiI (middle), and a merged image (right). *B. longum* (C) and *L. plantarum* (D) stained with DAPI (left), with CMO (middle), and a merged image (right).

**Figure 3.**
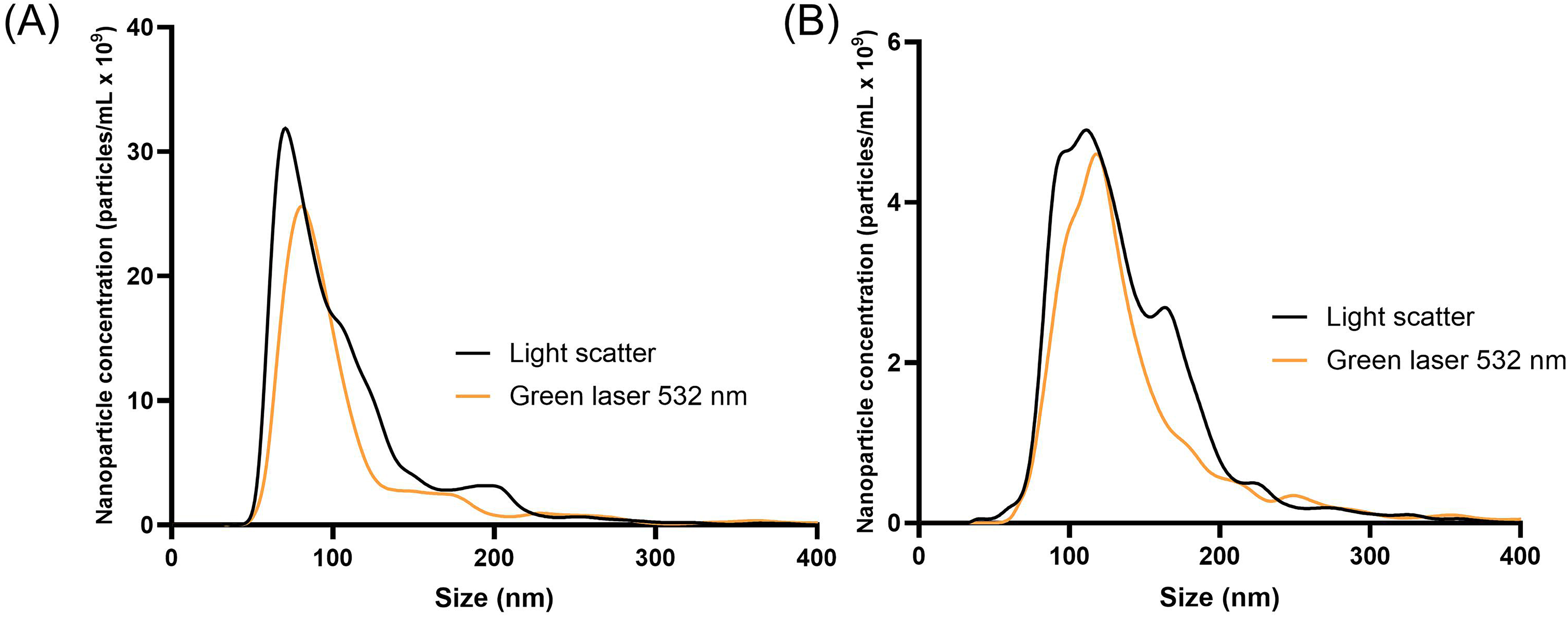
NTA detection of Bl-EVs and Lp-EVs after staining with CMO. CMO-stained nanoparticle detection of extracted and purified Bl-EVs (A) and Lp-EVs (B) using NTA with the light scatter (black lines) or the green laser 532 nm (orange lines) with the 565 nm long pass filter of the NanoSight NS300.

### 3.3. Vesiculogenesis of Bl-EVs and Lp-EVs using HR-TEM

Observations were carried out at the late exponential growth phase (Fig. 4). At the surface of *B. longum* and *L. plantarum*, membrane protrusions suggesting vesicle budding were observed (Fig 4A,B). This phenomenon was observed in all the examined cells, suggesting that it is not a random or sporadic event. However, differences in vesicle formation between the two species were observed. In *L. plantarum*, at an advanced stage of vesicle formation, an accumulation of organic matter was observed on the top of the emerging vesicles (Fig. 4D). This feature was observed for each emerging vesicle at the surface of *L. plantarum* while no similar biological process was identified during vesicle formation of *B. longum* (Fig 4C). Furthermore, differences in vesicle release between the two bacteria were also noted. For *B. longum*, vesicles appear embedded within the peptidoglycan layer before being released (Fig 4E) while for *L. plantarum*, visible disruptions of the cell wall surface in both sides of the emerged EVs were observed (Fig. 4F), suggesting that a wall-fracturing mechanism might facilitate Lp-EVs release. These findings indicate that the mechanisms of vesicle formation and release is likely different between the two Gram-positive bacteria.

**Figure 4.**
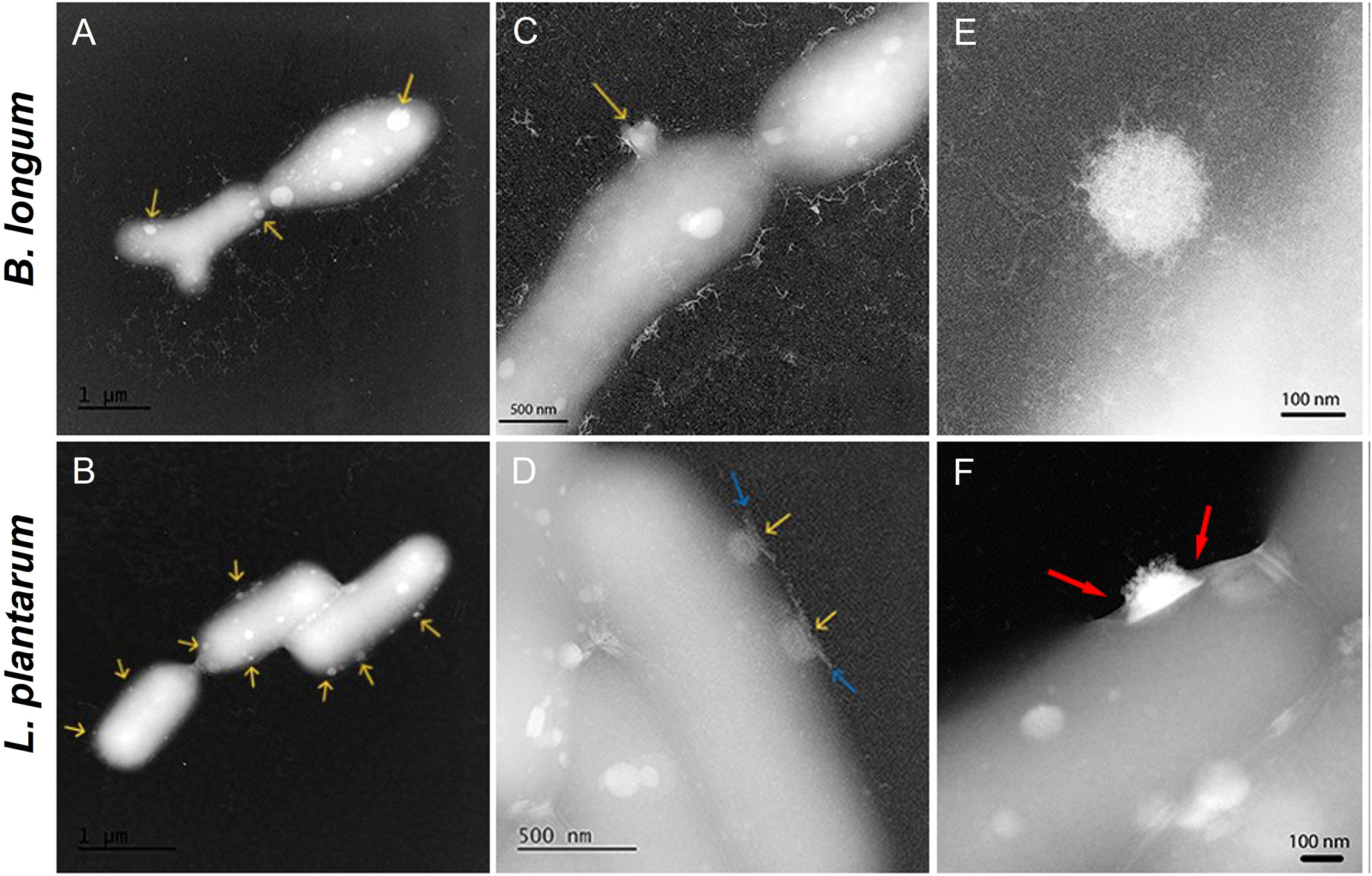
Vesiculogenesis of *B. longum* and *L. plantarum* as observed using TEM at 80 keV after negative staining. Yellow arrows indicate the location of BEVs at the surface, blue arrows show the biological matter accumulation during the Lp-EVs formation, and red arrows point out the membrane disruption before Lp-EVs release.

### 3.4. Impact of AMX exposure on Bl-EVs and Lp-EVs production rate

Bl-EVs and Lp-EVs were isolated from cultures in the presence of a sub-inhibitory concentration of AMX, previously determined by measuring the MIC for each strain. When *B. longum* was exposed to AMX, a significant increase of the Bl-EVs concentration was measured compared to the untreated control, irrespective to the bacterial growth (Fig. 5B). In opposite, the concentration of Lp-EVs was not significantly different for *L. plantarum* compared to the untreated control (Fig. 6A). However, vesicle size was not affected for Bl-EVs and Lp-EVs after antibiotic exposure (Fig. 5A). These results suggest that AMX can differentially modulate the production rate of bEVs but not their size, depending on the bacterial species.

**Figure 5.**
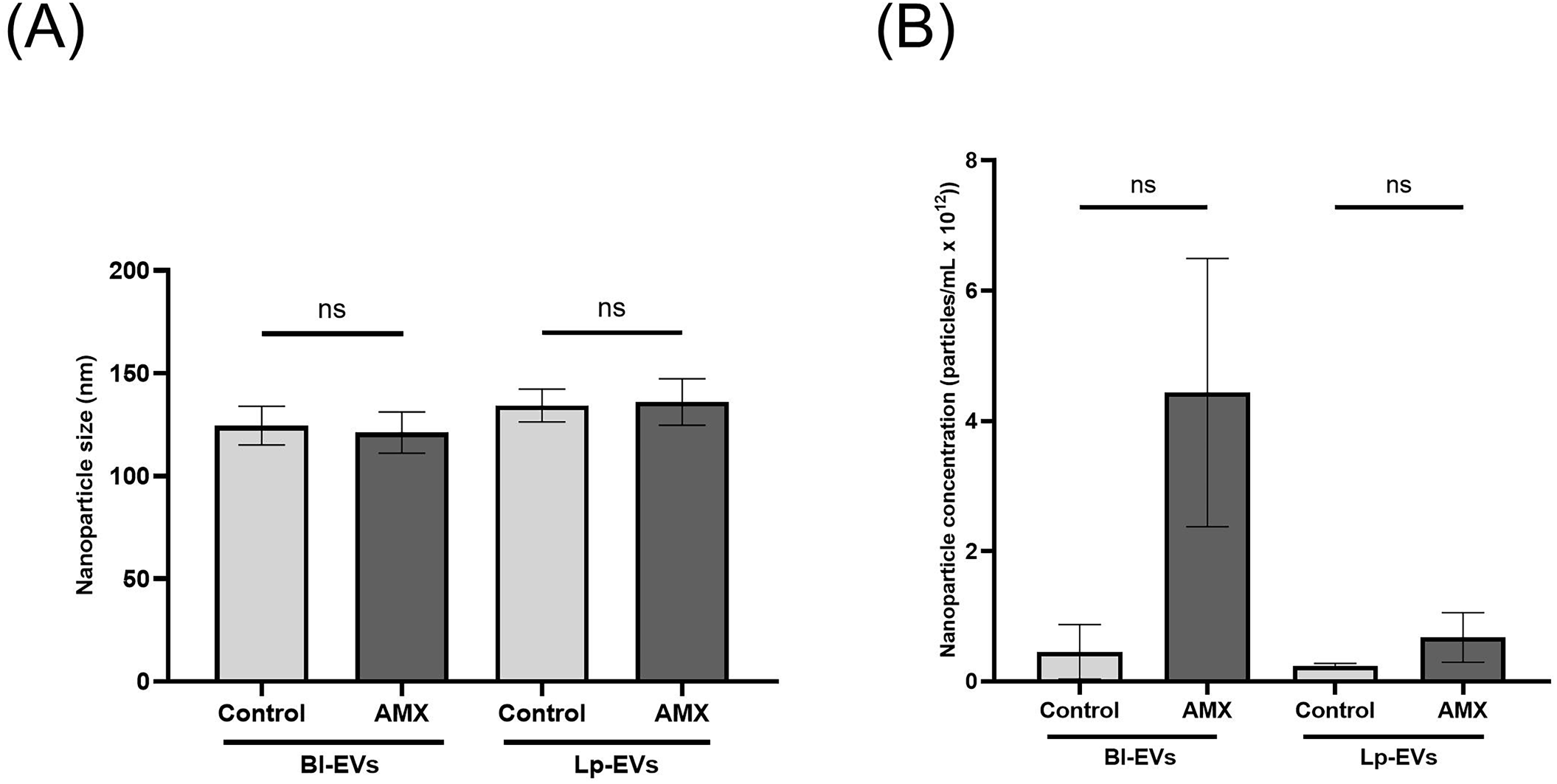
The bEVs concentrations of *B. longum* and *L. plantarum* under sub-inhibitory concentrations of amoxicillin. The bacterial concentration at the end of each culture was not significantly different among the three independent biological replicates (3.81 × 10^9^ ± 1.57 × 10^9^ cfu/mL for Bl and 2.69 × 10^9^ ± 3.77 × 10^8^ cfu/mL for Lp) and the cultures with AMX (4.27 × 10^9^ ± 2.01 × 10^9^ cfu/mL for Bl and 3.18 × 10^9^ ± 6.84 × 10^7^ cfu/mL for Lp). (A) Size distribution of bEVs under the same conditions measured by NTA. Mean of 3 biological replicates of nanoparticle concentrations according to their size. (B) Nanoparticle concentration measured by NTA in control and AMX-treated conditions (0.5 µg/mL for *B. longum* and 0.0625 µg/mL for *L. plantarum*. Statistical significance is indicated as ** *p*< 0.01, \*\*\**p*< 0.001 according to the One-way ANOVA test of the three independent replicates for each condition.

**Figure 6.**
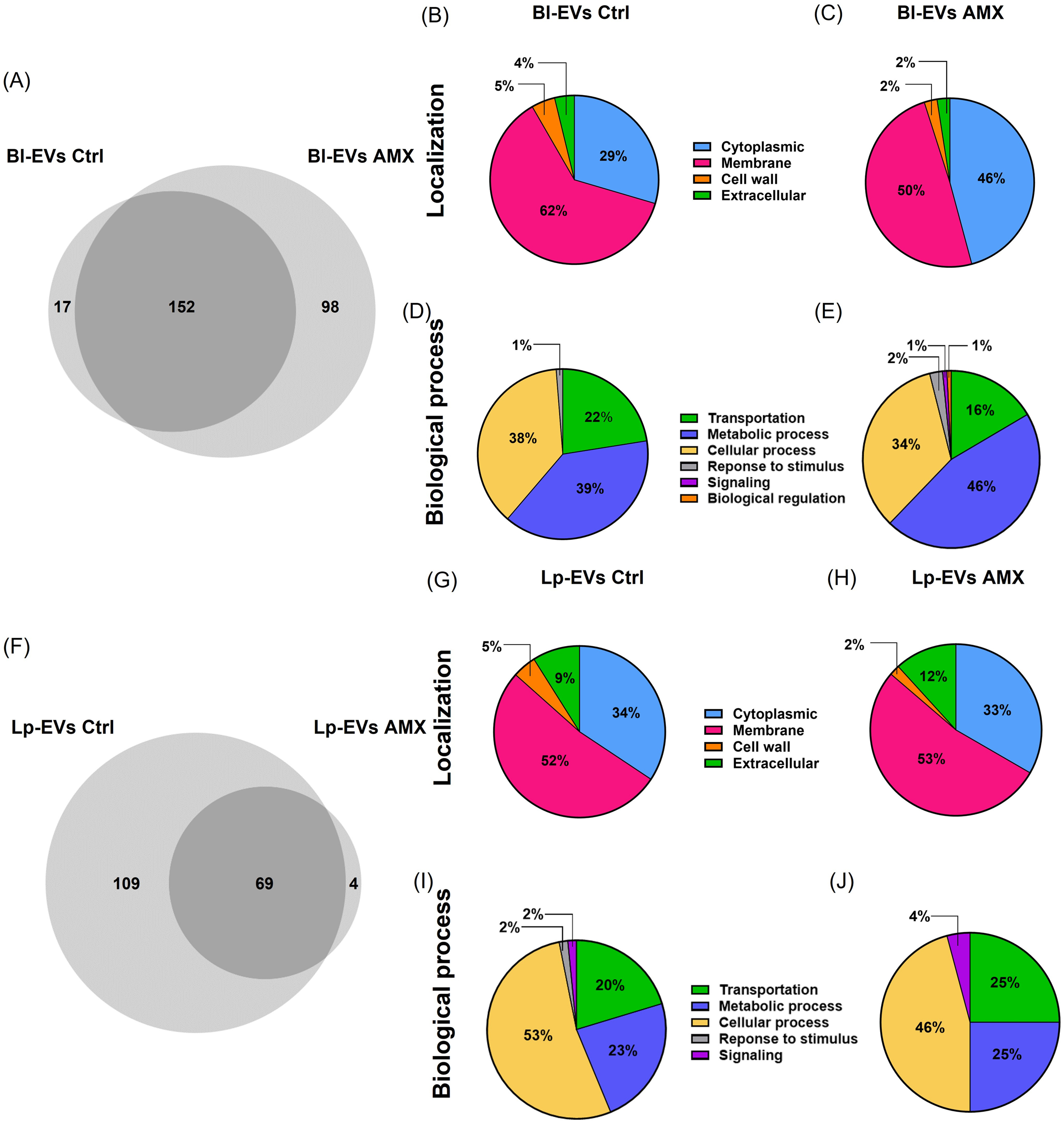
Proteovesiculomic analyses. Venn diagrams representing the number of common and different proteins present under control and AMX-treated conditions for Bl-EVs (A) and Lp-EVs (F) from three independent replicates for each condition. Bacterial Psortb-III localization predictions of proteins in Bl-EVs under control (B) and AMX-treated conditions (C). Bacterial Psortb-III localization predictions of proteins in Lp-EVs under control (G) and AMX-treated conditions (H). GO of main biological processes in Bl-EVs under control (D) and AMX-treated conditions (E). GO of main biological processes in Lp-EVs under control (I) and AMX-conditions (J).

### 3.5. Changes in vesicle protein fingerprintings after AMX exposure

Vesiculoproteomic profiling resulted in identifying a mean of approximatively 300 different proteins in Bl-EVs and Lp-EVs from cultures with or without AMX (Tab. S1, Fig. 6A,F). By comparing the common identified proteins among the biological replicates, 98 proteins in all replicates were newly detected in Bl-EVs after AMX-cell treatment while only 4 were newly detected in Lp-EVs (Tab S2). Correlatively, a drastic loss of cargo richness was observed in Lp-EVs as compared to the one in Bl-EVs after AMX-cell treatment. Predicted localization of proteins in Bl-EVs under control showed a predominance of membrane proteins, followed by cytoplasmic proteins and to a lesser extend proteins coming from cell wall and those predicted as extracellular proteins (Fig. 6B). After AMX-cell treatment, Bl-EVs were more enriched in cytoplasmic proteins while for Lp-EVs, localization prediction revealed no major shifts in protein distribution across compartments between the control and the AMX-treated conditions (Fig. 6G,H). The proteins detected in Bl-EVs and Pl-EVs come from mainly transport, metabolic, and cellular processes (Fig. 6D,E,I,J). In Bl-EVs, AMX exposure was associated with the presence of proteins related to the response to stimulus category, as well as the emergence of additional functional categories such as signaling and biological regulation (Fig. 6E). In contrast, in Lp-EVs proteins related to the response to stimulus category were not identified while proteins associated with signaling were identified after AMX-cell treatment (Fig. 6 J). Among the 98 common proteins detected in Bl-EVs after AMX exposure, 80 have been already detected in one or two independent biological replicates in the control conditions, indicating that these proteins became predominant in Bl-EVs after AMX exposure as they were detected in all biological replicates and 17 proteins were specifically detected in Bl-EVs in AMX conditions (Tab. I). For Lp-EVs, the four common proteins detected after AMX conditions have been already detected in one or two biological replicates in control conditions (Tab. 1). For Bl-EVs after antibiotic treatment, the specific identified proteins were mainly involved in membrane transport, metabolism, cell division, stress response, regulation, and quorum sensing. For Lp-EVs, the 4 proteins detected under AMX were associated with four distinct functions, including riboflavin metabolism, carbohydrate metabolism, extracellular protein, and membrane homeostasis (Tab. 1).

**Table 1.**
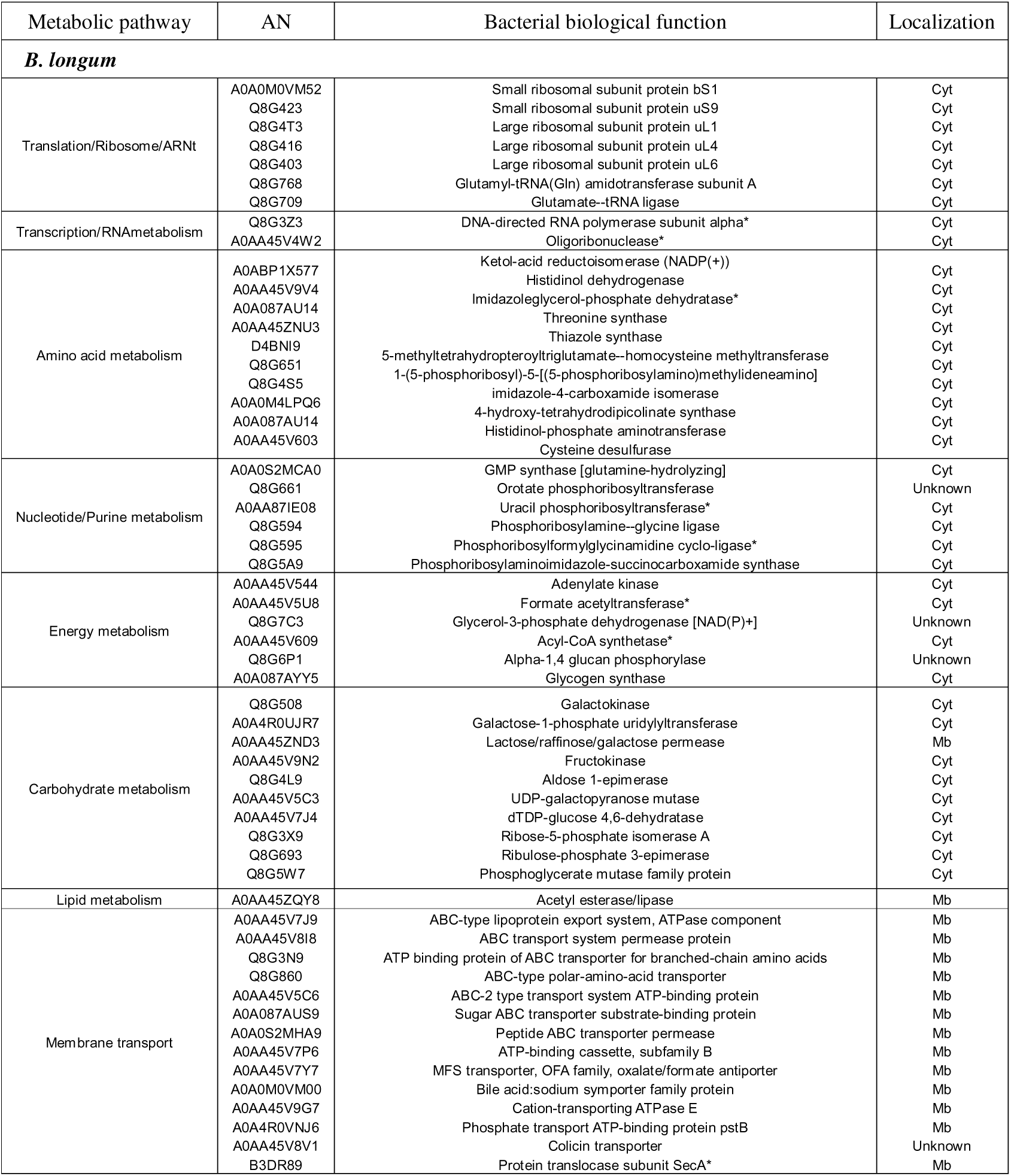
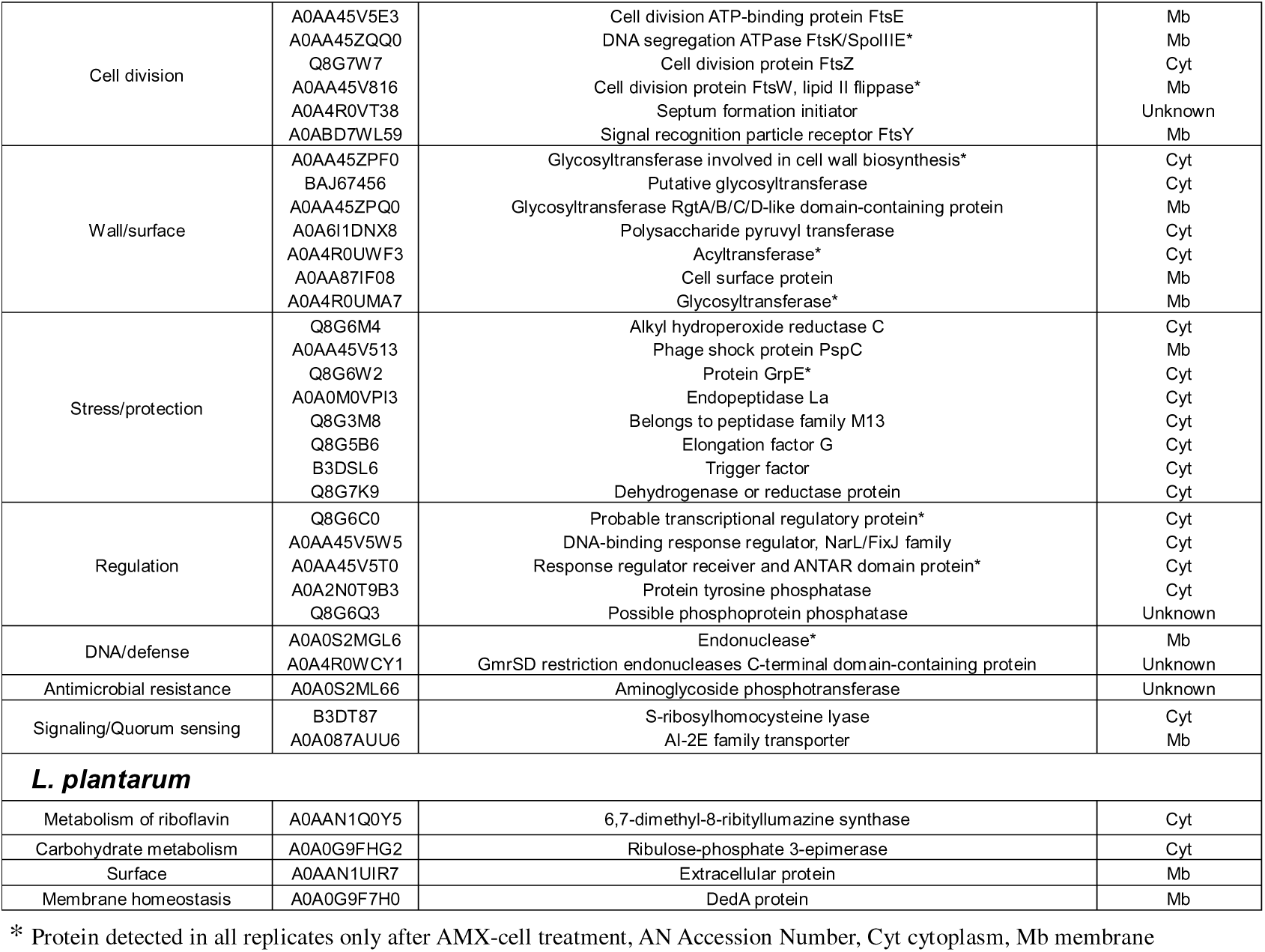
Unique biological functions acquired by Bl-EVs and Lp-EVs under AMX-treated conditions.

## DISCUSSION

Bacterial EVs, which act as vehicles for bioactive molecules, are recognized for their involvement in a wide range of biological processes, including intercellular communication and modulation of immune responses (Yi et al., 2025; Toyofuku et al., 2023). Gram positive bacteria with probiotic properties, including *Lacticaseibacillus paracasei*, *Propionibacterium freudenreichii*, *Lacticaseibacillus rhamnosus*, *Lactobacillus gasseri* and *Staphylococcus aureus*, have been reported to produce and release mEVs with anti inflammatory, infection protective, neuroprotective and immunomodulatory activities (Tartaglia et al., 2018; Ñahui Palomino et al., 2019; Choi et al., 2020; Rodovalho et al., 2020; Haas-Neill et al., 2022). In the early stages of life, the role of bEVs has not been yet documented, notably in the establishment of the symbiosis between gut primo-colonizing bacteria and the maturation of the main physiological systems during the first 1000 days of human life. In addition, the impact of antibiotherapies on first gut commensal bacteria have not been documented.

The isolation and characterization of bEVs remain a main methodological challenge due to the lack of a gold standard protocol (Hendrix et al., 2023). Although no universal method has been established, a consensus is emerging to harmonize sample collection, isolation, and analysis procedures in EVs research. In line with this, MISEV2023 guidelines have been regularly updated (Welsh et al., 2024). Following them, our results showed that a three-step protocol combining ultrafiltration, ultracentrifugation, and SEC efficiently isolated EVs from *B. longum*, and that this approach was extendable EVs of *L. plantarum*. The choice and combination of purification steps strongly influence bEVs quality and purity (Hendrix et al., 2023). In our workflow, centrifugation and clarification were used to remove bacterial cell and cellular debris, tangential ultrafiltration was used to concentrate vesicles and to limit contamination with small soluble molecules, ultracentrifugation was applied to separate EVs from >100 kDa soluble molecules while SEC was applied to purify vesicles. These steps ensured to perform studies on more purified vesicle fractions than those aiming at harvesting only enriched fractions such as those reported by Morishita *et al*., (2021), for instance. The elution profile obtained in this study is in line with optimized mEVs isolation protocol, where early SEC fractions contain vesicles and later fractions mainly contain soluble proteins (Watson et al., 2021). Furthermore, the absence of significant difference in concentration and size between F2 and F3 confirmed the homogeneity of the nanoparticle population extracted. The mean sizes ranging from 119 to 127 nm and the concentrations of approximately 2 × 10¹¹ nanoparticles/mL from *B. longum*, are consistent with previously reported ranges (Nishiyama et al., 2020; Mandelbaum et al., 2023). Likewise, the mean sizes (133–139 nm) and the concentrations of 2 × 10¹¹ nanoparticles/mL from *L. plantarum* fall within the expected range as obtained by Bajic et al., 2020 and Kim et al., 2022. These results demonstrate not only the reproducibility but also the cross-species applicability for Gram-positive bacteria. Thus, this protocol was validated to provide a reliable basis for future studies on mEVs composition and biological function. According to MISEV2023 guidelines, particles quantification by NTA or similar technology is essential to assess concentration and size distribution but cannot distinguish EVs from non-vesicular nanoparticles. Therefore, lipophilic fluorescent dyes were used as a complementary strategy to confirm vesicle membrane origin. Dyes such as DiI (Kim et al., 2016; da Silva et al., 2022), DiO (Champagne-Jorgensen et al., 2021), DiD (Jones et al., 2020), and PKH63 (Yang et al., 2022) are commonly used for bEVs detection. Unexpectedly, in our study, membrane of single bacterial cells of *L. plantarum* could not be all detected using DiI. However, CMO, usually used to label eukaryotic-derived EVs (Carnell-Morris et al., 2017) was more efficient to detect all DAPI-positive cells in both strains. Differences likely arise from the physicochemical properties of the dyes and the presence of a denser peptidoglycan layer in Gram-positive bacteria than in Gram-negative ones. In addition, the DiI, a highly hydrophobic carbocyanine, can form micelles and produce nonspecific signals (Chen et al., 2023), while CMO is amphiphilic and integrates efficiently into lipid bilayers. The used of fluorescence-coupled NTA with CMO-labeled particles revealed that most of the scatter-light-NTA detected nanoparticles are CMO-positive ones, confirming the mEVs detection for both strains. Altogether, these results provide the first demonstration of the CMO efficiency for the labeling of Gram-positive derived EVs.

In Gram-negative bacteria, vesicle biogenesis generates OMVs while lytic processes release IOMVs and EOMVs (Toyofuku et al., 2023). In Gram-positive bacteria, vesicle formation was once considered unlikely. In our study, with to the use of HR-TEM observations, the EVs release process seemed to be different between *L. plantarum* and *B. longum*. For the former strain, local wall perturbations and accumulation of organic material at budding sites were observed, consistent with a transient peptidoglycan rupture. This aligns with mechanisms described in *Bacillus subtilis*, where prophage-derived endolysins locally perforate the cell wall (Toyofuku et al., 2017). The structural organization of lipoteichoic acids (LTA), wall teichoic acids (WTA), and the autolysin Acm2, which regulates cell wall remodeling (Palumbo et al., 2006), may contribute to this process. Although further investigations have to be performed to elucidate EVs formation and release by *L. plantarum*, our observations confirmed the hypothesis of localized peptidoglycan disruption. According to our observations of the latter strain, Bl-EVs appeared to integrate progressively into the peptidoglycan layer before being released. This different way of release could be due to the unique structure of the lipoglycans of *Bifidobacteria*. While *L. plantarum* displays a typical lactic acid bacteria cell wall architecture, *B. longum* possesses a peptidoglycan matrix which is densely coated with lipoglycans composed of galactofuranan chains substituted with glycerophosphate and anchored to the cell membrane via a glycolipid, rather than with polyglycerol phosphate chains substituted with sugars, resulting in a thicker, more polysaccharide rich and structurally distinct cell wall (Balaguer et al., 2021). These findings would suggest that Gram-positive EVs biogenesis is different according to species with the limitations of this study applied on only the type-strain for each species. Hypothetical release mechanisms for EVs generated by Gram-positives include (i) turgor-driven extrusion; (ii) localized peptidoglycan degradation; and (iii) deformation of vesicles through pores smaller than their diameter (Brown et al., 2015). However, further analyses are required to better understand the shaping wall cell architecture, the enzymatic remodeling pathways, and the membrane composition during EVs release.

Amoxicillin (AMX) is one of the most commonly prescribed antibiotics during infancy, mainly for acute otitis media, bronchiolitis, pharyngitis, colds and urinary tract infections (CDC, 2024; Brustad et al., 2025). In a study covering 39,971 newborns, 44.3% of them received at least one course of antibiotics, with 78.5% of them starting their first course within the first three days of life. Among newborn treated, amoxicillin was prescribed in 53.1% of cases (Martin-Mons et al., 2021). In newborn treated with a 7 day course of AMX for acute respiratory infection, total bifidobacterial counts were preserved but the composition and diversity of the *Bifidobacteria* were markedly altered (Mangin et al., 2010) with a complete disappearance of *B. adolescentis* species, a significant decrease of *B. bifidum* but a stable amount of *B. longum* and *B. pseudocatenulatum/B. catenulatum*. A stable occurrence of *B. longum* was also reported in 5-days-treated adults with AMX as compared to the decrease of the gut bifidobacterial species (Mangin et al., 2012). In opposite, *Lactobacilli* dominated the bacterial community in controls, but its relative abundance was significantly reduced following AMX treatment (Graversen et al., 2020).

In our study, the exposure to a sub-inhibitory concentration of AMX induced a significant increase of Bl-EVs production without altering the size distribution while Lp-EVs production remains unchanged. This dissociated response pinpointed that bEVs production could be affected by antibiotic exposure, depending on the Gram-positive strain. In *S. aureus,* β-lactams including, ampicillin, oxacillin, imipenem, cefotaxime and cefoxitin induced a dose-dependent rise in bEVs release (Kim et al., 2020a; Huang et al., 2025). Similar bEVs upregulation has been described in group B *Streptococcus* exposed to ampicillin (Pell et al., 2025). Amoxicillin, as a beta-lactam as well, targets and inhibits cell wall synthesis by covalent binding the penicillin-binding proteins which are enzymes involved in the synthesis of the terminal step of peptidoglycan cross-linking in both Gram-negative and Gram-positive bacteria (Bush and Bradford, 2016). Inhibiting peptidoglycan transpeptidation could have been related to an increase of Bl-EVs release. However, the exposure to other classes of antibiotics including macrolides, fluoroquinolones, or aminosides could also impact the bEVs yield. For instance, sub-inhibitory concentration of gentamicin increases bEVs release in *S. epidermidis*, (Zaborowska et al., 2020). In addition, Gram-negative derived EVs are also enhanced in the presence of sub-inhibitory concentrations of metronidazole or levofloxacin for *Helicobacter pylori*, and the exposure to ciprofloxacin and polymyxin B for *Escherichia coli* (Bauwens et al., 2017; Krzyżek et al., 2025). The production of bEVs can also be modulated by the growth environment, including pH, temperature, oxidative stress and antimicrobial peptides. In addition, bEVs production could also vary between strains of the same species as demonstrated on *H. pylori* strains (Krzyżek et al., 2025). On another hand, studies also reported no increase vesiculation after an antibiotic treatment as for *L. plantarum* under AMX in our study. For instance, in *S. aureus*, vancomycin, erythromycin or kanamycin do not increase vesicle production compared with controls (Kim et al., 2020a; Huang et al., 2025). In *H. pylori*, clarithromycin does not significantly alter vesicle production. Altogether, the production of bEVs under antibiotic pressure seems not be to related to Gram-staining features, neither to the antibiotic mode of action including binding to ribosomes for macrolides, binding to DNA for fluoroquinolones or peptidoglycan transpeptidation for b-lactams. As the vesicle release seems to be different between *B. longum* and *L. plantarum*, further analyses should focus on EVs release and regulation molecular mechanisms. In addition, a better understanding of the EVs release regulation would pinpoint whether the formation of EVs inside the cell or the EVs cellular release process is affected by the regulation.

Nonetheless, most of the reported studies concerned EVs released by bacterial pathogens. Indeed, commensal and symbiont bacteria could be also affected by antibiotics. Then, our results extend the research on beneficial bacteria and more specifically on gut primo-colonizing bacteria such as *B. longum* and *L. plantarum*. The higher production of Bl-EVs under AMX indicated that bacteria-host signaling in the early stages of life might be altered by antibiotherapies. Vesiculoproteomic profiling indicated that approximately 300 statistically-validated proteins are transported by mEVs. A wide variability in the number of proteins detected in EVs from beneficial bacteria, ranging from a few dozen to more than one thousand, has been reported (Krzyżek et al., 2023). However, proteovesiculome studies have detected about one hundred proteins, supporting the relevance of our findings (Harrison et al., 2021; Rodovalho et al., 2021). Moreover, regardless of the condition, most of the proteins identified in our vesicles were predominantly membrane-associated. A similar diversity enrichment in membrane proteins has also been described in the EVs from *L. plantarum* BGAN8, in which 91.5% of the proteins specifically detected in EVs were membrane-localized (Bajic et al., 2020). This finding is consistent with the presumed biogenesis and release of bEVs through the cell wall (Brown et al., 2015). Beyond their cellular localization, the proteins identified in Bl-EVs and Lp-EVs were mainly associated with cellular process, metabolic process and transportation. For example, in Bl-EVs, the identified proteins are mostly involved in functional categories corresponding to metabolic pathways, ribosomes, and secondary metabolite biosynthesis, followed by ABC transporters (Mandelbaum et al., 2023). Under AMX exposure, additional functional categories emerged in Bl-EVs, including signaling and biological regulation. Similarly, signaling-related proteins were also observed in Lp-EVs, whereas the response to stimulus category disappeared in the presence of AMX. These results show that AMX exposure not only alters vesicle protein composition, but also reshapes their functional profile, which warrants a more detailed examination of the proteins specifically detected under this condition.

In Bl-EVs, AMX exposure led to a marked remodeling of the vesicular proteome, with the appearance of 98 proteins, mainly involved in membrane transport and carbohydrate and amino acid metabolism. The presence of transport and metabolic proteins suggests that these bEVs may contribute to the acquisition, processing, and delivery of nutrients from the surrounding environment in a bioavailable form, potentially benefiting both the bEVs and the host, particularly in nutrient-rich niches such as the gut (Krzyżek et al., 2023). Several proteins related to stress response and cellular protection identified in our study were also reported by Nishiyama et al., 2020 into EVs of *B. longum*. For instance, the presence of AhpC which is involved in oxidative stress response could detoxify peroxides, reactive oxygen species, and nitrogen- and sulfur-containing compounds (Zuo et al., 2014). We also identified GrpE, which belongs to the dnaK operon which participates in the general stress response in bifidobacteria (Ventura et al., 2005). Trigger factor likewise points to the activation of the chaperone system involved in the folding of newly synthesized proteins, in cooperation with DnaK, thereby limiting aggregate formation under stress (Merz et al., 2008). In addition, endopeptidase La indicates active protein quality control, likely required to remove damaged proteins under antibiotic stress (Kirthika et al., 2023). The identification of PspC further strengthens this interpretation, as this protein is associated with envelope stress and membrane integrity maintenance (Ravi et al., 2024). These findings indicate that, in the presence of AMX, *B. longum*-derived EVs might mobilize a coordinated bEVs response aimed at protecting EVs, preserving protein homeostasis, and maintaining envelope stability. Further analyses considering the protection of EVs from degradation would be fruitful in the understanding of the sustaining role of Bl-EVs in the protein protection from damage and degradation.

Several proteins involved in membrane transport, cell division, and cell wall remodeling were also detected in Bl-EVs in AMX conditions, including SecA, an essential ATPase of the Sec system responsible for protein translocation across the cytoplasmic membrane via SecYEG, as well as FtsZ, and FtsK/SpoIIIE, which are involved in cell division (Chen and Beckwith, 2001; Loose and Mitchison, 2014; Cranford-Smith and Huber, 2018; Chan et al., 2022). Their presence points to a substantial perturbation of the bacterial envelope and division dynamics, which is consistent with the mode of action of beta-lactams on peptidoglycan synthesis (Bush and Bradford, 2016). In this context, bEVs may reflect a structural and functional adaptation mechanism of *B. longum* to AMX. Several regulatory proteins were also detected in BL-EVs as the signaling-related factors such as LuxS and the AI-2 transporter, indicates that bEVs might contribute to the bacterial regulation and communication system. Indeed, LuxS is the enzyme responsible for the synthesis of the AI-2 precursor, a molecule involved in inter- and intraspecies communication (Turroni et al., 2025). In bifidobacteria, the luxS gene is conserved and strains are able to produce AI-2, supporting the existence of a functional signaling system in this genus (Yuan et al., 2008). In addition, the LuxS/AI-2 system has been associated with stress adaptation, biofilm formation, and, in lactic acid bacteria, tolerance to harsh intestinal conditions (Meng et al., 2022). In this context, the enrichment of these regulatory proteins in Bl-EVs in AMX conditions indicates that Bl-EVs potentially vehicle bioactive molecules involved the kingdom interplays. In the opposite, Lp-EVs showed a very reduce gain in proteins after AMX-cell treatment, suggesting that *L. plantarum* engages a more restrained response strategy than *B. longum.* Considering that *B. longum* is less affected by AMX in newborn gut microbiota (Mangin et al., 2010; Mangin et al., 2012) and that AMX stimulates Bl-EVs production, further analyses should explore the potential shift functions of EVs produced under antibiotic pressure. Moreover, this research pinpoints that although some bacterial population remain unchanged under antibiotic pressure, their potential of interaction and communication through EVs could be altered.

## CONCLUSIONS

In conclusion, this study establishes a robust workflow for the isolation, labeling and characterization of EVs produced by the early gut Gram-positive colonizers *B. longum* and *L. plantarum*. Our findings displayed a dissociated response of *B. longum* and *L. plantarum* for EVs release process, for EVs production under antibiotic pressure, and for EVs protein content shifting after antibiotic-cell treatment. This suggests that gut microbiota-derived bEVs might shift their functions under antibiotic treatment according to strains. Further researches should focus on the shifting content and function of EVs to better decipher how exposures and experiences in the early-stage of life could contribute to the alteration of the microbiota-host symbiotic interplay.

## Author contributions

**Anaïs Halbert** carried out all the experiments with the help of **Amandine Dupuy** for EVs characterization using NTA and wrote the initial draft of the manuscript. Protein identification using mass spectrometry was perfume by **Lisa Wallart** and **Julie Hardouin**. **Odile Tresse** conceived, designed and oversaw the study. **Odile Tresse** helped to optimize the protocols. AH performed the statistical and proteomics analyses. **Odile Tresse** and **Hervé Blottière** brought their expert critical point of view of the current work. All authors have critically reviewed the published version of the manuscript.

## Acknowledgments

The authors would like to thank Eric Gautron for the measurements performed using the IMN’s characterization platform PLASSMAT, Nantes, France and Emmanuelle Dé (UMR-CNRS PBS 6022, Rouen, France) for fruitful discussions about proteovesiculomic. Anaïs Halbert received a fellowship from Ecole Doctorale Biologie Santé, Pays de la Loire. This work was supported by the Nantes Institute of Digestive System Diseases and the SanteDige Foundation. The authors from Nantes (France) are part of the network Grand Ouest-EV.

## Data Availability

The MS raw data are available in Proteomics Identifications Database (Pride) repository under accession PXD078131.

## Data Availability statement

All data generated or analyzed during this study are included in this published article.

## Notes

https://www.ebi.ac.uk/pride/markdownpage/pridesubmissiontool

### Competing Interest Statement

The authors have declared no competing interest.

